# Erythrocytic plasma membrane calcium ATPase (PMCA4b) variations mediate redox imbalance to determine artemisinin sensitivity in malaria

**DOI:** 10.1101/2024.12.17.628803

**Authors:** Priya Agrohi, Swati Garg, Shreeja Biswas, Preeti Maurya, Vijay Kumar, Bidhan Goswami, Samir Kumar Sil, Mrigendra Pal Singh, Sunil Kumar Chand, Sneh Shalini, Om P Singh, Gunanidhi Dhangadamajhi, Sivaprakash Ramalingam, Prashant Kumar Mallick, Shailja Singh

## Abstract

Artemisinin resistance in *Plasmodium falciparum* remains a major public health challenge for the ongoing malaria elimination programs. Till now, artemisinin resistance is linked with the variations in the genome of *P. falciparum* which are mainly clustered in the Pfkelch13 gene. Several studies reported artemisinin resistance in the malaria endemic areas without finding any evidence of genetic variation in parasite for this occurrence. This study interrogated whether host genetic variations have any association with artemisinin sensitivity of malaria parasite. We investigated the relationship of 1) genetic variations in the human ATP2B4 gene and 2) the resultant surface expression of plasma membrane calcium ATPase (PMCA4b) with the intraerythrocytic levels of calcium, reactive oxygen species (ROS) and the resistance towards artemisinin in the intraerythrocytic parasite. This study found negative correlation of PMCA4b expression level with intraerythrocytic calcium and ROS levels. Further in-vitro growth assays revealed that artemisinin sensitivity is reduced in the parasites growing within the RBCs having low PMCA4b and high oxidative stress. Overall, this study highlights the strong association of host PMCA4b in cellular calcium mediated redox imbalance, which significantly contributes to artemisinin resistance in malaria parasite. Hence, this is the first study to document the effect of host variations on artemisinin sensitivity of the parasite. We further emphasize that many redox modulating RBC polymorphisms that are prevalent in malaria endemic areas could influence artemisinin resistance and can serve as potential biomarkers for predicting therapeutic response. Thus, detailed population specific research may provide new insights for personalized malaria treatment and may inform future drug development strategies.

**Significance:** The human host and malaria parasite share a closely interconnected relationship hence host variations could influence parasite drug resistance. This study demonstrates that reduced PMCA4b expression, the primary calcium efflux pump in RBCs, significantly increases the erythrocytic calcium and reactive oxygen species (ROS) levels, thereby decreasing the *P. falciparum* growth as well as artemisinin sensitivity. Many ROS inducing RBC polymorphisms, such as sickle cell, thalassemia, G6PD deficiency, PMCA4b variations etc. have emerged as strong malaria protective traits in several studies in malaria endemic areas. But these traits could also influence artemisinin resistance and can serve as potential biomarkers for predicting therapeutic response and highlights the importance of host erythrocytes oxidative microenvironment surveillance in monitoring antimalarial drug resistance.

## Introduction

Malaria elimination requires the concerted efforts of all the nations to achieve the goal of world free of malaria. However, the disease remains the major cause of child mortality worldwide, killing more than a million children in Africa alone each year even after introduction of various antimalarial therapies (1, 2). World Health Organization (WHO) has recently put forward a global technical strategy for malaria elimination by 2030 and recommends the stronger surveillance of antimalarial drug efficacy and resistance(3).

Malaria has co-evolved with human host since ages and has acted as one of the strongest selection forces on human genome (2). It is suggested that many human genetic polymorphisms are the result of selection for malaria resistance. For example, malaria endemic regions have witnessed the evolutionary selection of blood group antigens, hemoglobin chains and human lymphocyte antigen (HLA) classes which are protective of malaria (4), (5). Several erythrocyte defects that provide protection against malaria such as sickle-cell disease, thalassemia, glucose-6-phosphatase deficiency, etc. are prevalent in endemic areas (6). Alongside, introduction of antimalarial drugs due to human intervention is altering parasite population dynamics resulting in emergence of multi-drug-resistant genotypes (7). Presumably, when humans were evolving genetically to fight malaria the parasite was evolving to become resilient towards drug interventions.

Till now artemisinin-based therapies are the backbone of malaria treatment, due to their effectiveness against *Plasmodium falciparum* malaria. However, recent reports of artemisinin resistance (ART-R) have raised concerns about the sustainability of these therapies. ART-R is characterized by reduced killing of ring-stage *P. falciparum*, resulting in delayed parasite clearance (8). ART-R has been linked with the mutations in the Pfkelch13 protein propeller domain (9) however there are reports of artemisinin treatment failure without any mutations in K13 domain of the parasite (10, 11). Recently host erythrocyte reactive oxygen species (ROS) had also been linked with resistance of malaria parasite towards artemisinin(12). This report indicated that genetically variant host RBCs may also influence the sensitivity of antimalarial drug. Surprisingly, the effect of human host variations on the artemisinin resistance has not been investigated yet. Even a very recent study indicated the role of two other parasite encoded proteins, PfRAD5 and PfWD11, in ART-R but still misses out on the contribution of host in this phenomenon (13, 14).

Intraerythrocytic oxidative stress can be linked with many physiological factors including the intracellular calcium levels of RBCs (15–17) which is primarily regulated by the plasma membrane calcium ATPase (PMCA4b). PMCA4b is encoded by ATP2B4 gene which has emerged as strong malaria resistance candidate in several GWAS and case-control studies in various parts of the world. Variation in the regulatory region of ATP2B4 were found to have a very strong association with resistance to severe *P. falciparum* malaria in a recent GWA study conducted in Kenya, Ghana and across many sites in Africa, Asia, and Oceania(18–22). These variation presents in ATP2B4 gene decreased the likelihood of SM by 40% in African children while also reducing the odds of placental *Plasmodium falciparum* malaria in African women by 64% (23). These independent and diverse research findings strongly indicate the significant role of ATP2B4 gene in infection of malaria parasites to human host **(Appendix Table 1**). However, the role of PMCA4b mediated regulation of intracellular calcium in redox imbalance and antimalarial drug resistance has not been investigated yet.

**Table 1.**
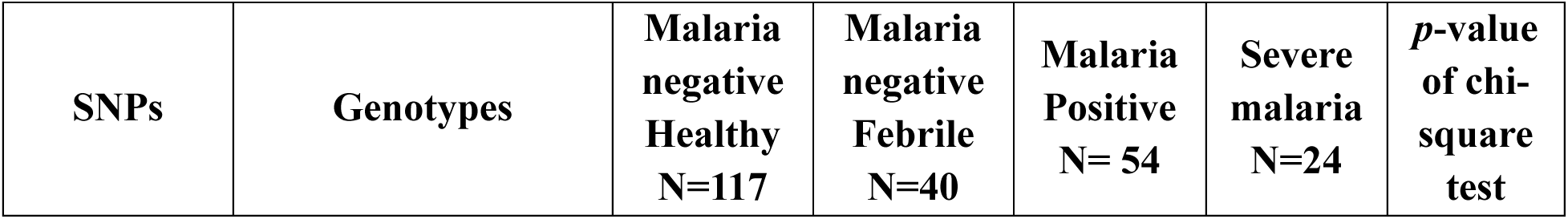
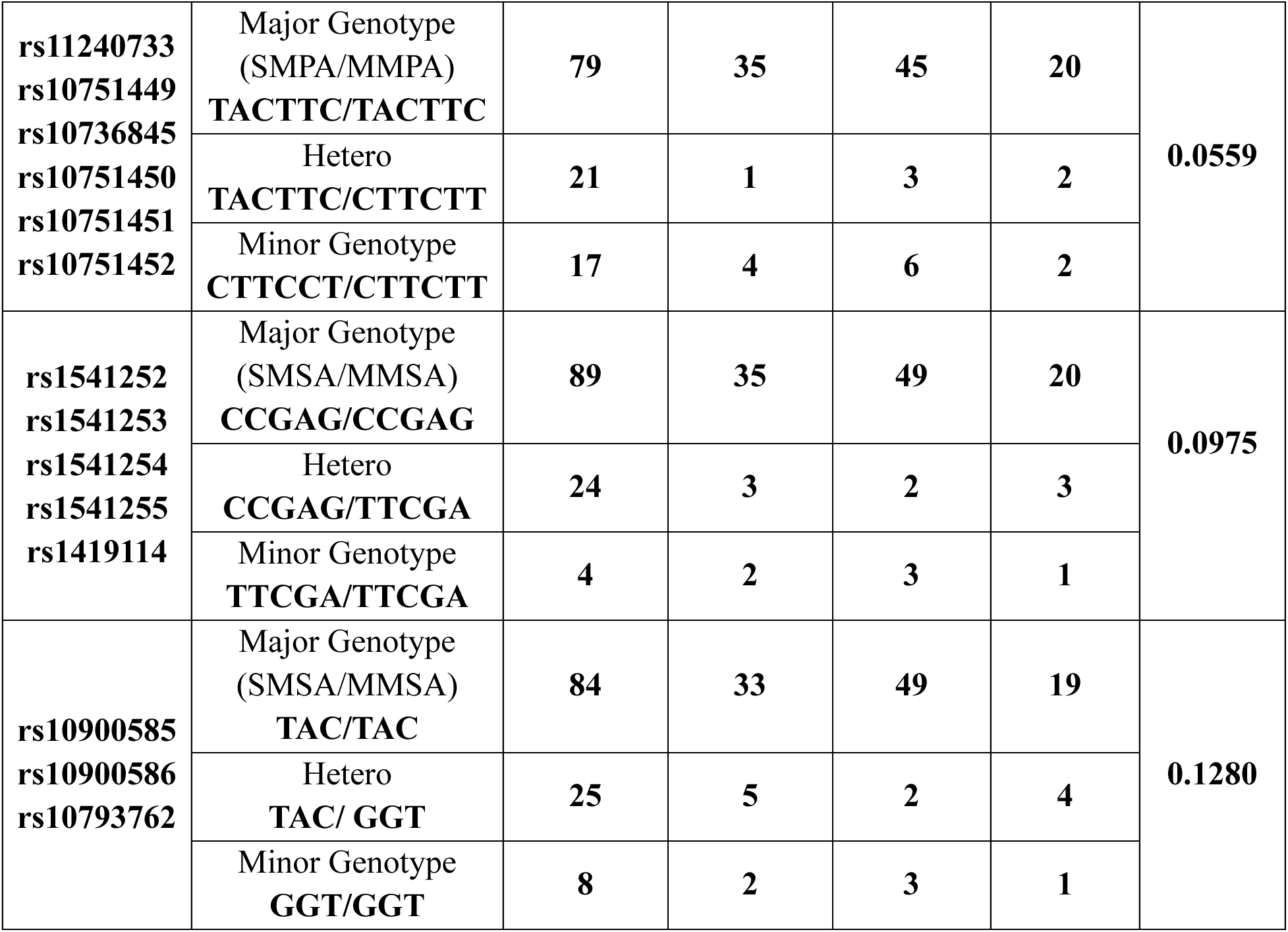
Prevalence of genotypes in Different Sample Groups. . This table displays the prevalence of major, minor, and heterozygous genotypes present in three different regulatory regions of the gene in different malaria outcome, including malaria positive, malaria negative, febrile but malaria negative and severe malaria of Indian population.

For the first time, this systematically designed experimental study investigated the relationship of human genetic variations in ATP2B4 gene with the resultant surface expression of PMCA4b, the intraerythrocytic levels of calcium, reactive oxygen species (ROS) and the resistance towards artemisinin in the intraerythrocytic parasite. We demonstrate that the protein expression of PMCA4b negatively correlates with the intraerythrocytic calcium and ROS levels. Further *in-vitro* growth assay results revealed that artemisinin efficacy is reduced in the parasites growing within the RBCs having low PMCA (high calcium and ROS). This study majorly highlights the role of host PMCA4b in conferring artemisinin resistance to the parasite. Several studies reported artemisinin resistance in the malaria endemic areas without finding any genetic evidence for this occurrence. Our data provide an explanation for this and advocates the need to consider the human factors while assessing this phenomenon. Thus, there is need for population specific research to understand the role of host genetic factors influencing artemisinin resistance in the malaria parasite.

## Results

### The complex nature of the PMCA4b encoding regulatory ATP2B4 genetic variations was not found to be associated with malaria susceptibility in the studied Indian population

Several SNPs of ATP2B4 gene have been studied for their association with malaria protection. This study focusses on SNPs selected on the basis of either strong GWAS signal or experimental research (**Appendix Table 1**). These SNPs are clustered in three regions of the gene: Enhancer [1^st^ intron], 5′UTR and Second intron (Figure 1a). According to the previous studies, alleles of these SNPs have been categorized as: SMPA (severe malaria protective allele), SMSA (severe malaria susceptible allele), MMPA (mild malaria protective allele), and MMSA (mild malaria susceptible allele). This categorization is based on their association with either severe/mild malaria protection or susceptibility (Figure 1a, SMPA, MMPA-illustrated by green color and SMSA, MMSA-illustrated by red color).

**Figure 1:**
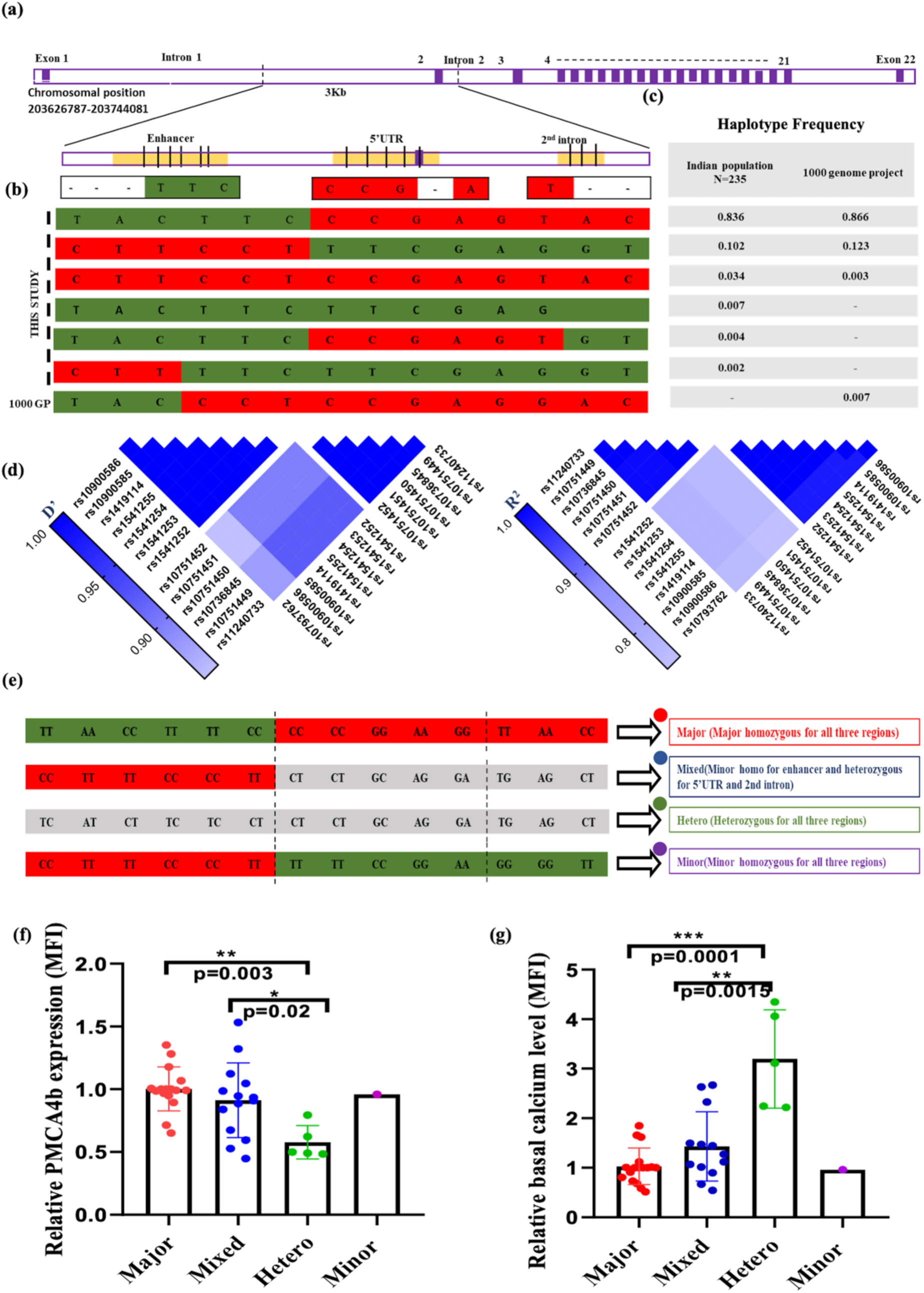
Characterization of ATP2B4 gene variations and their impact on PMCA4b expression and basal calcium levels in the studied Indian Population. **(a)** Schematic diagram of the ATP2B4 gene highlighting the three regulatory regions containing variations. A total of 14 variations were identified by sangar sequencing, with 8 previously reported mutations and 6 more mutations (represented by dash) described either protective (Green color) or susceptible (Red color). The zoomed image illustrates the specific locations of these mutations within the regulatory regions. **(b)** Distribution of haplotypes in the ATP2B4 gene among the studied Indian population and the 1000 Genomes Project. Green and red color represents the previously known protective and susceptible alleles. **(c)** Frequency of identified haplotypes within the studied population compared to global populations included in the 1000 Genomes Project. **(d)** Heatmap of linkage disequilibrium (LD) values (D’ and r²) across the 14 SNPs in the ATP2B4 gene. The color gradients indicate the degree of association (D’) and correlation (r²) between each pair of SNPs. **(e)** Representation of the four ATP2B4 genotypes observed in the Indian population, selected for further analysis. Each genotype is annotated in a color-coded box, with corresponding colors matched to those used in subsequent figures for easy reference. **(f)** PMCA4b expression levels and **(g)** Basal calcium levels across the four ATP2B4 genotypes. Both measurements are plotted with mean fluorescence intensity (MFI) on the y-axis and genotypes on the x-axis. *p*-values are indicated in the figures.

Using the UCSC genome browser, regulatory mechanisms like histone modification, DNase hypersensitive region, and remap density plot were explored for these regions. The first intronic region acts as cis-regulatory elements (cCREs), likely regulating PMCA expression. The 5′UTR shows promoter like activity and predicted to serve as a secondary promoter specific to erythrocytes. However the, second intronic region, despite being associated with malaria protection in GWAS, did not exhibit any clear involvement in regulatory mechanism (Appendix figure 1).

Variants of ATP2B4 gene were explored in the Indi Genomes (Indian genome database)(24), which shows the frequency of altered alleles for all SNPs ranges from 0.8 to 0.9 in the Indian population (Appendix Table 1). Genotyping in 117 random malaria negative apparently healthy individuals identified 14 SNPs in the same regulatory regions (Figure 1a). According to the hardy-Weinberg test result all the analyzed SNPs are in hardy-Weinberg equilibrium (Appendix Table 3). Genotyping of these variants shows allelic frequency data corroborate with Indian database. Further investigation was extended to various malaria outcomes in other studies to check the frequency of these variations in *P. falciparum* positive and severe malaria cases.

Genotyping of 14 SNPs present in regulatory regions of ATP2B4 gene were performed in severe malaria and mild malaria cases. 6 SNPs were observed in enhancer region yield 3 genotypes: TACTTC/TACTTC, CTTCCT/CTTCCT, and heterozygous for all. In the 5′UTR, 5 SNPs were observed which yield 3 different genotypes: CCGAG/CCGAG, TTCGA/TTCGA, and heterozygous for all. Similarly, in the 2^nd^ intron, 3 SNPs were observed and yield three genotypes: TAC/TAC, GGT/GGT, and heterozygous. All the observed genotypes of studied regulatory regions showed equivalent prevalence across different malaria outcomes. (Table 1).

### The Indian population exhibit distinct pattern of Linkage Disequilibrium compared to global population

All these 14 SNPs are in 3kb DNA fragment spanning the chromosome region chr1:203681464-203684970 (Figure 1a). Haplotype analysis of all 14 SNPs identified 4 haplotypes in LD-link (a web-based application analyzes the linkage pattern in populations of 1000genome project) using all population groups from the world. In total six haplotypes were observed in the studied Indian population with various frequencies. Every haplotype observed in the Indian population displayed a mosaic pattern of alleles that are previously reported as protective or susceptible against severe malaria. Likewise in major haplotype, previously reported SMPA/MMPA (indicated by green color) were observed for enhancer region while previously reported SMSA/MMSA (indicated by red color) were observed for 5′UTR and 2^nd^ intron (Figure 1b). This similar pattern is also observed in haplotypes created by the 1000 genome project database (Figure 1c).

LD plot of all the SNPs show two blocks of variants present in complete linkage in Indian population, one having only promoter region and other one with 5′UTR and 2^nd^ intron (Figure 1d). However, in contrast, the LD plot from the 1000 genomes project the same set of variants displayed complete linkage among all the SNPs, suggesting a different LD pattern across the Indian population (Appendix figure 2).

**Figure 2:**
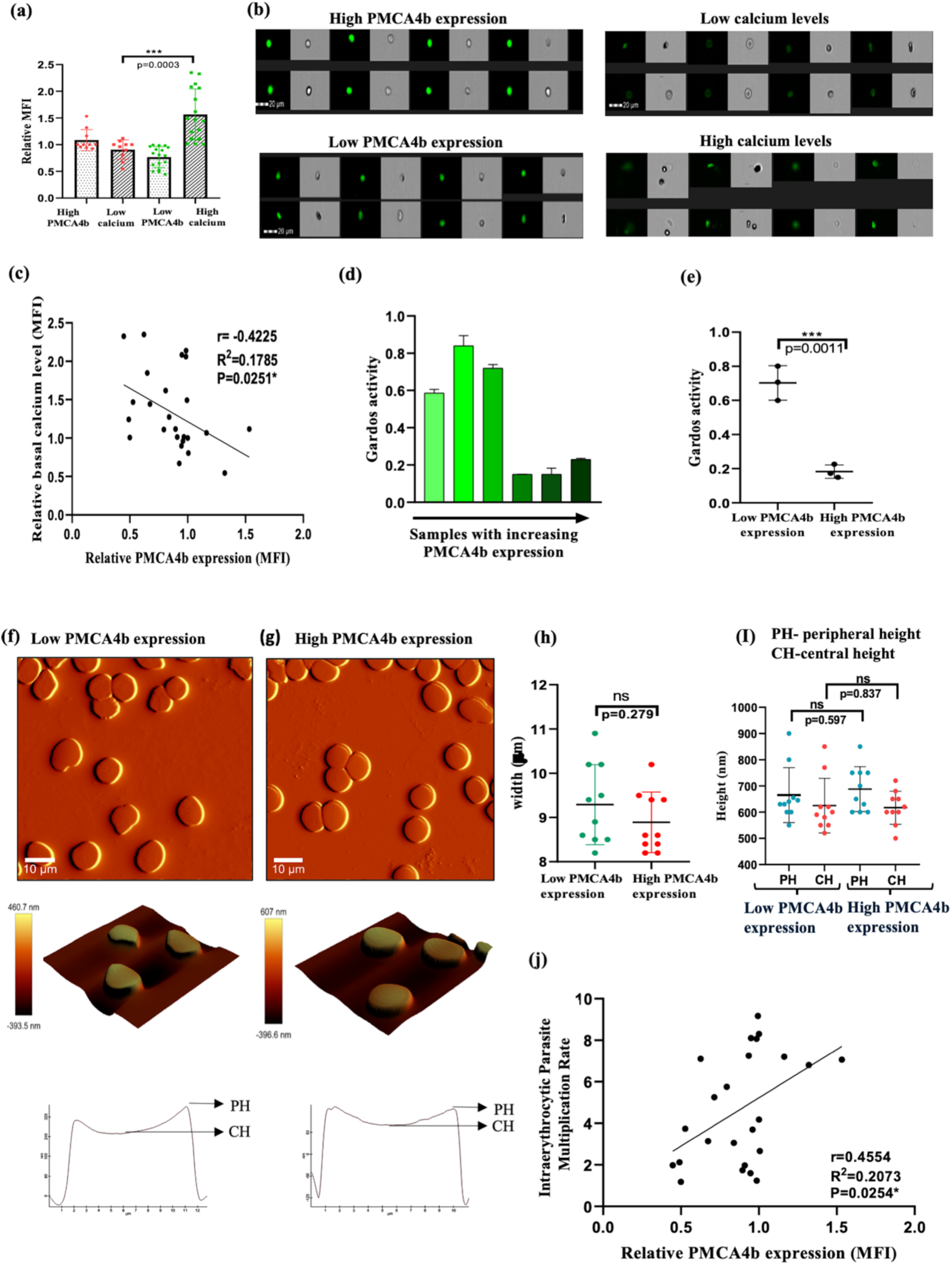
Impact of PMCA4b expression on Red Blood Cell morphology and structural measurements. **(a)** Stacked column graph depicting the relationship between high and low PMCA4b expression and basal calcium levels. **(b)** Representative images from imaging flow cytometry illustrating high and low PMCA4b expression and corresponding low and high calcium levels. **(c)** Scatter plot demonstrating the inverse correlation between PMCA4b expression and calcium concentration. Pearson correlation coefficient and *p*-values are indicated in the figure. **(d)** Measurement of Gardos channel activity in RBCs with different PMCA4b expression (expression in increasing order on x-axis) assessed using the FluxOR™ Potassium Ion Channel Assay. Fluorescence intensity was measured before stimulus which is considered as basal level and after stimulus. stimulation with thallium (Tl) is plotted on the y-axis, indicating potassium flux through the Gardos channel. **(e)** Combined activity of the Gardos channel in RBCs with high and low PMCCA4b expression levels. *p*-value indicating significance is indicated. **(f)** Atomic Force Microscopy (AFM) images of RBC with low PMCA4b expression. Scale=10 µm scale **(g)** AFM image of RBC with high PMCA4b expression. Scale=10 µm scale, illustrating surface characteristics and overall morphology reflective of the physiological state associated with PMCA4b levels. Zoomed AFM image (scale=6 µm), showcasing detailed structural features including roughness and height profiles that highlight the microtopography of RBCs with PMCA4b levels. **(h)** Width and **(i)** height measurements of red blood cells (RBCs) with respect to PMCA4b levels is plotted. **(j)** Scatter plot showing the direct relationship between PMCA4b expression levels and the growth rate of the *P. falciparum* 3D7 strain. Lower PMCA4b levels are associated with decreased parasite growth. Pearson correlation coefficient and *p*-values are indicated in the figure.

### Correlation study ATP2B4 polymorphisms with PMCA4b expression and basal calcium levels in RBCs

In studied population, a total of four genotypes were observed, representing combination of alleles for all the 14 SNPs (Figure 1e): Major, Mixed, Heterozygous and Minor. The effect of these genotypes on PMCA4b expression and basal calcium level were investigated. The graph (Figure 1f) shows relative PMCA4b expression levels across four genotypes. The data indicates significant differences in PMCA expression among these genotypes by using one way ANOVA test (*p*=-0.004). Further Tukey′s multiple comparisons test shows that heterozygous genotypes has significantly lower PMCA4b expression as compared to major (*p*=0.003) and mixed genotype (*p*=0.02). Basal calcium level was also investigated among these four genotypes (Figure 1g). one way ANOVA test indicates significant difference in calcium level among these genotypes (*p*=0.0001). Further Tukey′s multiple comparisons test shows that heterozygous genotypes has significantly higher basal calcium level as compared to major (*p*=0.0001) and mixed genotype (*p*=0.0015). the minor genotype displays a higher PMCA4b expression level and lower calcium level, though no statistical comparison can be conducted because of only single sample of it.

### Inverse association of PMCA4b Expression with intraerythrocytic Calcium Levels

To investigate the relationship between PMCA4b expression and basal calcium level, samples were assigned into two categories: high PMCA4b expression and low PMCA4b expression, the average basal calcium level in each group was compared which revealed significant difference between two categories (Figure 2a). Representative imaging flow cytometry images also illustrate high calcium staining in low PMCA expressing cells and vice versa indicating the reverse correlation (Figure 2b).

Following this observation, we analyzed the linear correlation of PMCA4b expression with intraerythrocytic calcium levels and linear regression test was performed among the samples irrespective of their genotypes. The regression analysis reveals an inverse correlation between PMCA expression and intracellular basal calcium level in RBCs (negative slop of −0.8664, 95% CI). Additionally, Pearson’s correlation coefficient also supports this correlation (*p*=0.0251) (Figure 2c). This observation suggests that as PMCA4b expression inversely correlate with basal calcium levels in RBCs.

### PMCA4b expression negatively regulates the potassium ion efflux Gardos channel activity

Elevation of intraerythrocytic calcium levels activate Gardos channels to efflux potassium ions outside RBCs. Gardos channel activation was investigated by FluxOR™ potassium ion channel assay. This assay measures fluorescence intensity, indicating potassium flux through the Gardos channel. Gardos channel activation was assessed at basal levels (without stimulus) and after stimulation with thallium (Tl). The fluorescence intensity of each sample was normalized with an average baseline fluorescence intensity measured before adding stimulus (TI). The result shows that RBCs with high PMCA expression (lower basal calcium level) exhibit less activity of the Gardos channel compared to RBCs with low PMCA expression (Figure 2d). The activity of the Gardos channel is directly influenced by basal calcium levels, with fluorescence intensity decreasing as basal calcium levels decreases (Figure 2e). The difference was statistically significant (*p*=0.0011) indicating the role of PMCA4b expression in regulation of RBC physiology.

Activation of Gardos channels facilitate echinocytosis of RBCs. To observe echinocytic RBCs under variable intraerythrocytic calcium levels, side scatter (SSC) analysis was performed using flow cytometry. SSC values correspond to cell granularity and hydration status, with increasing SSC of erythrocytes suggest cell dehydration. Pearson’s correlation test indicates a positive trend between basal calcium level and SSC values; however, this correlation was not statistically significant (p=0.0695) (Appendix figure 3), suggesting no association of PMCA4b derived calcium with RBC morphology.

**Figure 3:**
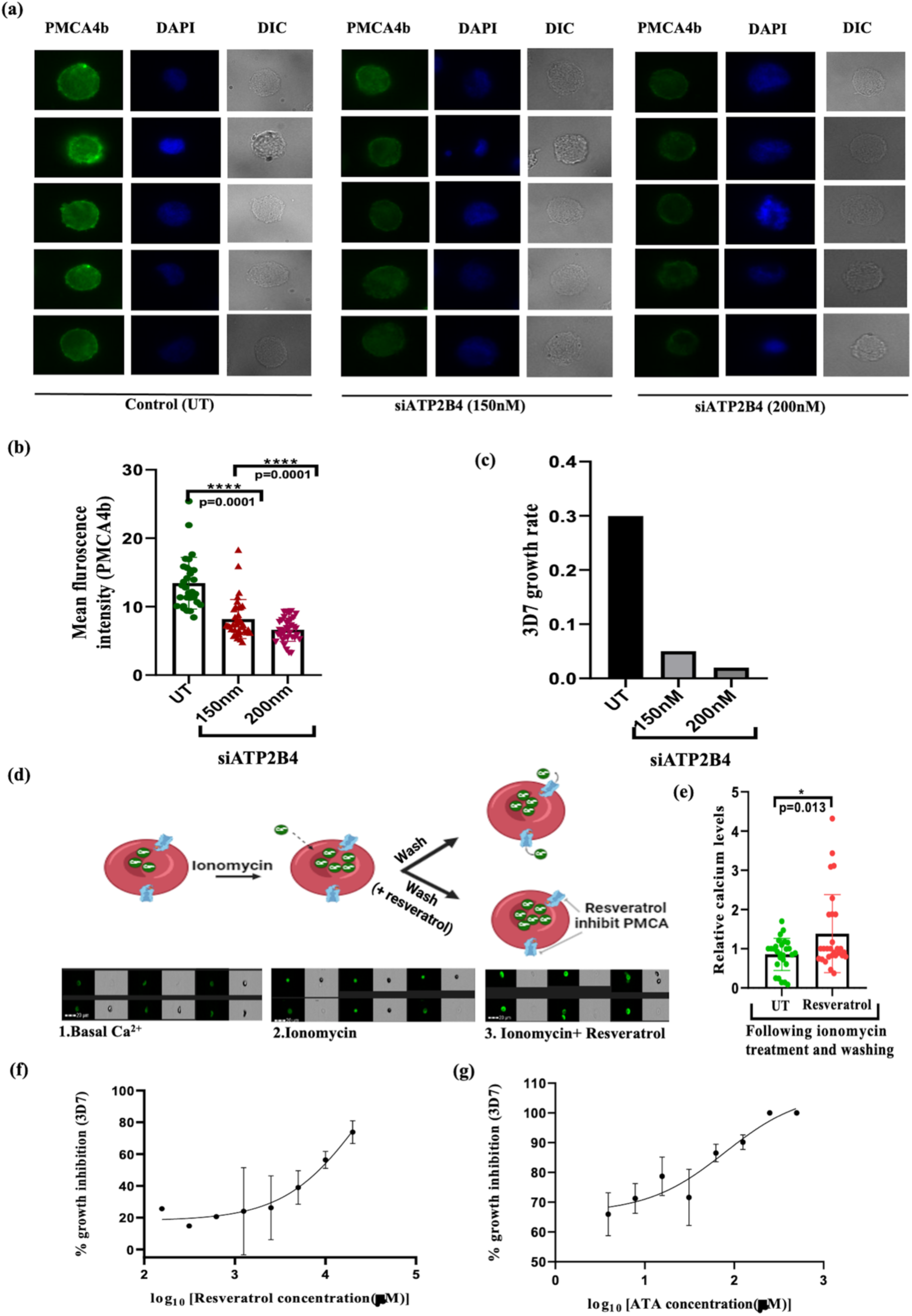
Resveratrol and siRNA mediated PMCA inhibition: Effects on calcium levels, ROS production, and *P. falciparum* growth in donor RBCs and BEL-A cells. (a) siRNA mediated silencing of ATP2B4 gene in BEL-A cells (Bristol Erythroid Line Adult cells). Representative fluorescence microscopy images showing PMCA4b expression analyzed by fluorescence microscopy. (b) Mean fluorescence intensity (MFI) analysis of PMCA expression in untreated (UT), 150 nM siRNA, and 200 nM siRNA-treated cells, indicating effective reduction of PMCA expression with increasing siRNA concentration. (c) *P. falciparum* growth rate in UT and siRNA-treated cells (150 nM and 200 nM), showing a potential impact of PMCA downregulation on parasite growth. (d) Schematic diagram of the calcium analysis in RBCs, along with representative fluorescence images. RBCs were treated with ionomycin in calcium-containing medium to induce calcium influx, and basal fluorescence was measured. Following a 2-hour treatment with resveratrol (50µM), fluorescence was measured again to assess the effect of PMCA inhibition on calcium efflux. (e) Effect of resveratrol on intracellular calcium levels in RBCs. The observed fluorescence intensity in ionomycin serve as basal calcium level and fluorescence in RBC with ionomycin + resveratrol reflects changes in calcium levels after PMCA inhibition. *p*-values are indicated in the figure. (f) Effect of PMCA4b inhibition by resveratrol and (g) ATA on parasite growth. Growth Inhibition Assay (GIA) curves show the antimalarial effects of resveratrol and ATA on parasite growth after 48 hours of treatment.

We performed Atomic Force Microscopy (AFM) to examine in detailed the morphological and structural changes in RBC with high and low PMCA level. 2D and 3D AFM images of RBCs with low (Figure 2f) and high (Figure 2g) PMCA expression shows no significant changes in surface features. In addition to lack of visual morphological differences, quantitative measurement of RBC width and height were also analyzed to determine if PMCA level influence the RBCs size. Width (Figure 2h) and height (Figure 2i) results shows that RBCs with low PMCA expression had slightly decreased height compared to those with high PMCA expression, though these changes were not statistically significant (p=0.38 for peripheral height, p=0.36 for central height). Similarly, there is no statistically significant difference in the width of RBCs with high and low PMCA level (p=0.185).

### Expression of PMCA4b on erythrocytes positively associates with *P. falciparum* Growth

Previous studies correlated ATP2B4 SNPs with malaria protection. Hence, the effect of PMCA4b expression on intraerythrocytic growth of *P. falciparum* was monitored. Briefly purified schizonts were incubated with RBCs having variable PMCA4b expression and parasite growth was monitored after 48h. The result shows a significant positive relationship between PMCA4b expression and growth rate of malaria parasite indicating low PMCA4b expression may confer protection from malaria (Figure 2j).

### Downregulation of ATP2B4 expression by siRNA mimics the natural PMCA4b polymorphism for parasite growth phenotype

To confirm that reduced PMCA4b expression is an important factor in parasite growth, ATP2B4 mRNA was downregulated using siRNA. A decrease in PMCA4b expression was observed following transfection of 150 nM and 200 nM of ATP2B4 targeting siRNA (Figure 3a, 3b). Statistical analysis using one-way ANOVA with Tukey’s multiple comparisons test revealed a significant difference among the groups (P < 0.0001). Specifically, PMCA expression was significantly lower in cells treated with 150 nM siRNA (mean MFI = 8.18, p=0.0001) and 200 nM siRNA (mean MFI = 6.64, p=0.0001) compared to the untreated control group (mean MFI = 13.44). However, the difference between the 150 nM and 200 nM siRNA groups was not statistically significant (P = 0.0632). Additionally, shows that the inhibition of PMCA4b via siRNA at both 150 nM and 200 nM concentrations corresponded to a notable reduction in the growth rate of *Plasmodium falciparum* 3D7 (Figure 3c), confirming the previous observation that lower PMCA4b levels negatively impact parasite growth within RBCs.

### Chemical inhibition of PMCA4b activity by Resveratrol and ATA restricts parasite growth

Previous results indicate that low PMCA4b expression correlates with increased calcium level and reduced *Plasmodium falciparum* growth. To further validate this observation, intraerythrocytic calcium levels and parasite growth were monitored following inhibition of PMCA4b. To check the effect of resveratrol mediated PMCA inhibition on RBCs calcium level, donor RBCs were treated with ionomycin to increase the intracellular calcium followed by resveratrol treatment that will inhibit the efflux of calcium by PMCA4b inhibition (Figure 3d). Comparative analysis of calcium levels was performed in the presence and absence of resveratrol. Calcium levels were significantly high in resveratrol treated RBCs indicating inhibition of calcium efflux (mean difference: 0.5276 ± 0.2069; 95% CI: 0.1126 to 0.9425). An unpaired t-test confirmed this difference to be significant (P = 0.0137) (Figure 3e). Representative flow cytometery images illustrate basal calcium levels, calcium after ionomycin treatment, and calcium levels after treatment with ionomycin and resveratrol. Furthermore, resveratrol demonstrated dose dependent growth inhibition of *P. falciparum* parasites with an IC50 of 18.35µM (Figure 3f). Another inhibitor of PMCA4b, Aurintricarboxylic acid (ATA) was also tested for parasite growth inhibition that demonstrated an IC50 of 78.8µM (Figure 3g). These results suggest that inhibition of PMCA4b restricts parasite growth underscoring its potential as druggable target. This data is similar to the parasite growth in field samples with variable PMCA4b expression.

### Host PMCA4b associated redox imbalance alters parasite sensitivity towards artemisinin

High cellular calcium is often linked with high ROS levels. We further investigated the relationship between calcium and oxidative stress inside the RBCs using DCFDA dye as an indicator of ROS (reactive oxygen species). Linear regression analysis shows a significant positive correlation between calcium and ROS level. Indicating that increased calcium level correlates with higher ROS level. Further Pearson’s correlation coefficient confirms this correlation to be significant (p=0.0005) (Figure 4a).

**Figure 4:**
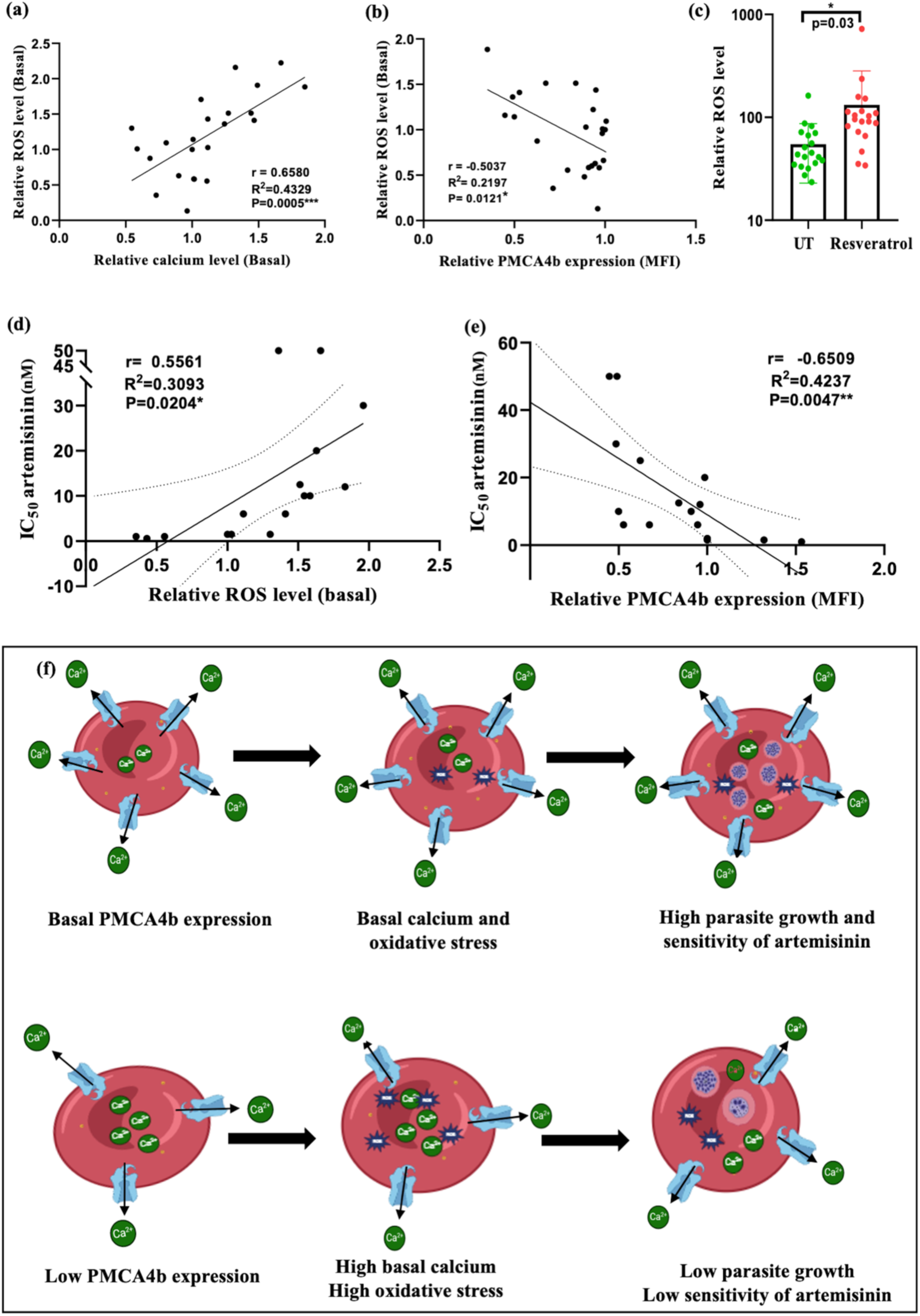
Role of oxidative stress of RBCs and PMCA4b expression in modulating Artemisinin Efficacy in *P. falciparum*. **(a)** Scatter plot illustrating a direct relationship between ROS and calcium levels in RBCs, suggesting that elevated calcium levels correlate with higher ROS production. **(b)** Scatter plot showing the inverse relationship between PMCA4b expression and ROS levels in RBCs, measured using DCFDA dye. Lower PMCA4b levels correlate with higher ROS production. **(c)** Effect of resveratrol on ROS levels in RBCs. Following a 2-hour treatment with resveratrol (50µM), cells were stained with DCFDA dye, and ROS levels were measured using flow cytometery. **(d)** Scatter plot illustrating the inverse relationship between ROS levels and artemisinin sensitivity, measured as IC₅₀ of DHA. **(e)** Scatter plot showing the direct relationship between PMCA4b expression levels and artemisinin sensitivity (IC₅₀). Pearson correlation coefficient and *p*-values are indicated in the figure. **(f)** Final hypothesis model showing the relation between RBCs PMCA4b expression, calcium levels, oxidative stress and artemisinin sensitivity.

Similarly, the inverse relationship between PMCA4b level and ROS level was observed. Pearson correlation coefficient confirms this negative correlation (p=0.0121) (Figure 4b) suggesting that reduced PMCA4b expression level leads to increasing calcium concentration which contributes in increased oxidative stress in RBCs. To further confirm that the change in ROS levels was due to PMCA4b polymorphism but not any other variation, the impact of resveratrol treatment on ROS production in RBCs was investigated. A paired t-test comparing basal ROS levels and ROS level following resveratrol treatment indicates a statistically significant increase in ROS generation after resveratrol exposure (P = 0.0355). The mean difference in ROS levels between the two conditions was 77.53, with a 95% confidence interval of 5.861 to 149.2 (Figure 4c) indicating the specific role of PMCA4b in redox imbalance.

Oxidative stress of host erythrocytes predisposes the malaria parasites to develop tolerance against oxidative damage causing drugs such as artemisinin. The correlation analysis between the ROS levels of RBCs and antimalarial IC_50_ of artemisinin shows a positive slope, indicating the loss of artemisinin sensitivity in the presence of redox imbalance. The regression analysis was statistically significant with a *p*-value of 0.0204 (F=6.717), suggesting that the slope is significantly different from zero. The correlation coefficient (r) of 0.5561 (95% CI: 0.1030 to 0.8181) supports a positive association, with an R-squared value of 0.3093. These results suggest that increases in the ROS level of RBCs are associated with higher artemisinin IC_50_ values (Figure 4d).

The corresponding relationship between PMCA4b expression levels of RBCs and *P. falciparum* sensitivity towards artemisinin was also investigated. A significant inverse correlation was observed between PMCA4b level and IC_50_ of artemisinin, as represented by the linear regression analysis, indicated that higher lower PMCA4b levels in RBCs were associated with increased IC_50_ of artemisinin in the parasite. The slope of the regression line was −33.35 (95% CI: −54.75 to - 11.94), significantly different from zero (F = 11.03, P = 0.0047), suggesting that variations in PMCA4b level have a measurable impact on artemisinin efficacy. Pearson’s correlation coefficient (r = −0.6509) further confirms a significant negative correlation, with a 95% confidence interval of −0.8619 to −0.2478 (Figure 4e). These data propose a new concept of host mediated evolution of drug resistance particularly in the areas where several forms of ROS inducing hemoglobinopathies are prevalent.

## Discussion

Artemisinin based combination therapies are the frontrunners in the malaria treatment and played major role in malaria elimination programme. Emergence of resistance in parasite against artemisinin or other partner drugs caused a serious threat in the global malaria elimination efforts. The reduced sensitivity of *P. falciparum* to artemisinin is caused by the mutations in the propeller domain of the kelch13 gene (PfK13) inducing dormancy and reduced metabolic activity in the parasites until the serum drug concentration is reduced. The active surveillance of mutations in K13 locus is being done to track artemisinin resistance. However, reports of delayed parasite clearance have also been reported without PfK13 mutations indicating the role of other factors in development of artemisinin resistance(10, 25). Recent studies also indicated role of WD40 and RAD5 in addition to K13 locus in delayed parasite clearance. (26, 27).

The parasite demonstrates an immense relationship with its human host; hence, the parasite biology cannot be studied independent of the host physiology. A number of studies indicated the role of host proteins such as Band 3, Glycophorins and Basigin as a crucial factor for parasite infection (28, 29). The polymorphisms in human PIEZO1, a mechanosensitive channel, are associated with malaria protection (30). Host Gardos channel inhibition have potent antimalarial activity indicating its role in disease pathogenesis(31). At the genetic level, multiple single nucleotide polymorphisms (SNPs) in the regulatory regions of ATP2B4 confer protection against severe and mild malaria (18, 19, 21–23, 32). Mostly, the studies were focused on African populations often overlooking the southeast Asian populations where both *P. falciparum* and *P. vivax* malaria are endemic.

This study was rationally designed to correlate both field and experimental data. It identified that the previously reported SNPs in ATP2B4 regulatory region are also common in Indian population but the protection against severe/mild malaria cannot be seen in limited samples. This could be attributed to different genetic makeup of Indian population from African population where most of the ATP2B4 studies were conducted suggesting that population-based host genetic differences have a huge influence on malaria protection. Additionally, ATP2B4 genotype does not associate strongly with the PMCA4b expression and basal calcium levels across the studied population. Thus, the protective effect of ATP2B4 SNPs, previously hypothesized to be due to the reduced expression of PMCA4b, could not be confirmed experimentally in the studied population indicating the involvement of additional regulatory mechanisms.

The PMCA4b expression strongly correlated with basal intraerythrocytic calcium levels and *P. falciparum* growth. Individuals with lower PMCA4b expression exhibit higher basal calcium and reduced parasite growth compared to those with higher PMCA4b expression. The high calcium mediated Gardos channel activation might cause Gardos effect leading to efflux of potassium ions. However, detailed analyses by AFM did not reveal significant structural or morphological changes in RBCs. This suggests that naturally occurring variations in PMCA4b expression might affect RBC physiology but their impact may not result in detectable alterations in RBC morphology under the normal conditions. Additionally high calcium was found to be positively correlated with erythrocytic ROS levels as reported in previous studies for nucleated cells(15). Thus, the malaria protective mechanism of RBCs with lower PMCA4b expression could not be attributed to changes in RBC morphology. Rather changes in intraerythrocytic calcium and ROS levels could be a determining factor in the malaria protective mechanism of RBCs with variable PMCA4b expression.

The role of PMCA4b expression in *P. falciparum* growth was further supported by Resveratrol-mediated inhibition of PMCA4b in RBCs and siRNA-mediated knockdown in BEL-A cells. Resveratrol increased intracellular calcium and demonstrated potent antimalarial activity (33). Similarly, in BEL-A cells, silencing of PMCA4b significantly suppresses parasite growth, highlighting its critical role in facilitating *P. falciparum* survival. These findings underscore the therapeutic potential of targeting PMCA4b as a strategy to combat malaria.

The study by Pires et al., identified that parasite survival mechanism to oxidative stress is similar to the artemisinin stress thus *P. falciparum* parasites growing under oxidative stress conditions can adapt and become less sensitive to artemisinin (12). Intraerythrocytic ROS levels exhibit a strong positive correlation with calcium levels and negative correlation with PMCA4b expression, suggesting that RBCs with low PMCA4b expression and high calcium concentrations exhibit elevated ROS levels, creating a cellular environment that could influence parasite response towards artemisinin. This hypothesis is further supported by growth inhibition assays, which demonstrate that parasites growing in RBCs with lower PMCA4b level and higher calcium level become less sensitive to artemisinin, as indicated by an increase in IC50 values with increasing ROS levels and decreasing PMCA4b expression. These findings highlight the intricate interplay between RBCs calcium homeostasis and oxidative stress in determining drug sensitivity of parasite surviving in such conditions hence emphasizing the potential need to account for RBC physiological states when optimizing antimalarial treatments.

Malaria endemic areas often see selection of host genotypes such as hemoglobinopathies, G6PD deficiency, which naturally have high ROS levels demonstrate some level of protection from malaria due to reduced parasite infection(34). But, once infected, parasite within them will be tolerant to ROS and hence to ROS inducing drugs such as artemisinin as well. Thus, evolution of host is predisposing them from some amount of malaria protection but making them highly vulnerable for drug resistance. Therefore, we propose that the screening of host redox levels is crucial for planning containment and elimination strategies of drug-resistant malaria. Furthermore, the parasites demonstrate partial host mediated drug resistance in our study suggesting that doses of artemisinin administered can be modulated in individuals susceptible for drug resistant malaria. To conclude, this study highlights the critical role of host polymorphisms in modulating *P. falciparum* growth and sensitivity towards artemisinin. Lower PMCA4b expression is associated with reduced parasite growth; however, once the parasite survives under conditions of elevated ROS, it becomes more adaptive and exhibit decreased sensitivity to artemisinin. These findings suggest that while evaluating artemisinin resistance, we should not only focus on parasite variability but also consider the broader spectrum of interactions with human host factors, such as variations in erythrocyte environments. This comprehensive approach could provide deeper insights into artemisinin resistance mechanisms and inform more effective intervention strategies.

## Methods

### 1. Ethics statement

The clinical samples used in this study, including those from individuals with severe malaria, febrile malaria negative, and uncomplicated *P. falciparum* malaria, were collected from previous studies. Blood samples from malaria negative apparently healthy individuals were collected after receiving ethical approval from the Institutional Ethics Committee of ICMR-NIMR, New Delhi, India. Written informed consent was obtained from all participants prior to sample collection.

### 2. Study population and sample collection

A total of 235 sample were used in this study, 117 blood sample were collected from healthy volunteers at NIMR Delhi in the form of DBS [dried blood spot] by finger pricking for DNA isolation. DNA samples from Febrile malaria negative (n=40) (manuscript in preparation), Uncomplicated *Plasmodium falciparum* malaria (n=54) (manuscript in preparation) and Severe malaria cases (n=24) (35) were taken from other studies. For other experiments 2 mL veinous blood was collected from the malaria negative apparently healthy donors.

### 3. Selection of SNPs and annotation of selected polymorphism

Many SNPs associated with malaria resistance have been identified in previous studies. While these SNPs are predominantly located in regulatory and non-coding regions spanning the first to third exons of the gene, this study prioritizes those with robust GWAS signals or replicated results from experimental studies (**Appendix Table 1**). UCSC Genome Browser (36) was used to analyze their functional significance and potential regulatory roles.

### 4. DNA isolation, PCR amplification and genotyping

Genomic DNA isolation was performed on dried blood spots (DBS) using the QiaAmp DNA minikit following the manufacturer’s instructions [Qiagen, Hilden, Germany]. Three sets of primers were designed using Primer-BLAST software to amplify the 3 kb regulatory DNA sequence of the ATP2B4 gene, including the enhancer region, 5′ untranslated region (5′UTR), and 2nd intronic region. Initial PCR amplification involved a 3 kb DNA fragment using forward primers for the enhancer region and reverse primers for the 2nd intron. A nested PCR was subsequently performed to amplify all three regions (**Appendix Table 2**). Nested PCR of 20 μL reaction volume used 1 μL of template DNA from the primary PCR. Some samples displayed variability in the amplification of the enhancer region, which required adjustments in primer sets [F 5′-AAAAGGCTGATGTGTGGCAG-3′, R 5′-CGGCACCAAGTTTTGTTCTGA-3′]. PCR products were electrophoresed on a 2% agarose gel [Promega/Ameresco] in 0.5 TBE buffer and visualized under UV transillumination at 302 nm using a gel documentation system. The PCR products (887 bp, 749 bp, and 603 bp) were cleaned using ExoSAP-IT™ [Thermo Fisher Scientific] and sequenced with forward and reverse primers using ABI BigDye Terminator v3.2 [Thermo Fisher Scientific]. The sequences were aligned and compared with the reference human genome GRCh38 [Genome Reference Consortium Human Reference 38] and analyzed for nucleotide variations using Mega 11.0 software(37), followed by evaluations of genotype and allele frequency across different groups.

The distribution of genotypes (Major, Minor, and Heterozygous) for each regulatory regions (Enhancer, 5′UTR and 2^nd^ intron) across the four malaria outcome groups was analyzed using Chi-square test for independence. The test assessed whether the distribution of genotypes was independent of the malaria outcome groups or if there were significant differences in genotype prevalence across groups. All analyses were performed with a 95% confidence interval and *p*-value less than 0.05 was considered statistically significant.

### 5. HWE, Sequence Phasing and Linkage Disequilibrium

Hardy-Weinberg equilibrium (HWE) analysis was performed based on allele frequencies to assess whether all alleles were in equilibrium. For linkage disequilibrium analysis, DNA sequence data were phased using DNAsp software (version 6). Phasing was performed for accurately inferring the haplotype phase of the diploid sequences, essential for proper estimation of haplotype diversity and other population genetic parameters. The phasing process utilized algorithms from HAPAR(39), PHASE, and fast PHASE. The level of linkage disequilibrium (LD) within the samples was measured using R² and D′ statistics, and the total number of haplotypes and segregating sites were determined using DNAsp software. Pairwise linkage disequilibrium for each genetic variation was computed and LD plots were generated using TASSEL(42) and GraphPad Prism 8.0.1.

### 6. PMCA4b protein expression and localization on RBCs

PMCA4b protein expression was measured by flow cytometry-based assay. In brief, 10-20 µL washed RBCs from each donor with different ATP2B4 genotype was fixed with 2.5% glutaraldehyde/PBS for 30 min at 4°C, permeabilized with 0.01% triton X100/PBS for 10 min at room temperature (RT) and blocked with 3% BSA in PBS. Permeabilized RBCs were then incubated with Anti-PMCA4b clone JA3 primary antibody (Merck Millipore) in 1%BSA/PBS for 1 hr at RT followed by secondary antibody, Alexa Fluor 488 goat labelled anti-mouse antibody (Life Technologies) in 1% BSA in PBS for 1h at RT. After antibody labeling, thin blood smear was made for the localization study and same sample were acquired using AMNIS imaging flow cytometer (Cytek AMNIS FlowSight, Manufacturer: Luminex Corporation, Austin, TX, USA). The results were analyzed using AMNIS IDEAS software, version 6.4, to determined MFI. Based on the imaging data, single cell population was selected excluding the doublets, clumps, debris and mean fluorescence intensity was measured. Thin blood smear slide was examined using a confocal microscope (Olympus, Shinjuku, Tokyo, Japan) with 100x objective for localization study.

### 7. Intracellular basal calcium concentration and Calcium efflux assay

Intracellular calcium concentration was assessed in the RBCs from different ATP2B4 genotypes with the help of Fluo-4AM calcium dye assay (43) with some modification. In brief washed RBCs from each donor was stained with 1 µM Fluo-4-AM dye (Molecular Probes/Invitrogen) for 30 min at 37°C in dark. After incubation, dye loaded RBCs were washed with 1XPBS and analyzed with imaging flow cytometry to assess the intracellular basal calcium concentration. To measure the activity of PMCA4b, Fluo-4AM stained RBC were loaded with calcium using 10 µM ionomycin ionophore (Molecular Probes/Invitrogen cat. 12,422) in incomplete RPMI medium containing 0.42 mM calcium. Calcium efflux from the RBCs were measure by flow cytometry at 0 minutes and 15 min following calcium loading. Effect of resveratrol (PMCA inhibitor) on calcium efflux was also assessed by using 50 µM resveratrol.

### 8. Measurement of intraerythrocytic ROS levels

Intracellular ROS (Reactive oxygen species) of RBCs was measured by DCFH-DA (2′,7′-Dichlorodihydrofluorescein diacetate) (D6883, Sigma-Aldrich) oxidation-sensitive fluorescent dye, by flowcytometry. DCFH-DA is non-fluorescence probe which can diffuse inside the RBCs where it deacetylated by esterase, producing DCF (Dichlorofluorescein), which emits fluorescence when oxidized by ROS. Washed RBCs were loaded with 10 µM DCFDA dye and incubated in dark for 40 minutes, after incubation RBCs were washed with 1X PBS and acquired in flow cytometry. Effect of PMCA inhibition or increased calcium concentration inside RBCs on ROS production was measured by treating RBCs with 50 µM resveratrol for 1 hour before staining with DCFH-DA dye.

### 9. Erythrocytic Flux OR™ Potassium Ion Channel Gardos activity Assay

The activity of Gardos channel in the RBCs with different PMCA4b expression was measured by thallium based FluxOR™ Potassium Ion Channel Assay (molecular probes, Life technology). In brief the iRPMI media was removed from the washed RBCs then RBCs were incubated in Flux OR II reagent dye in loading buffer (contains FluxOR™ reagent, a thallium sensitive fluorogenic indicator dye) for an hour in dark. it was followed by washing and 30 min of exposure to assay buffer (contains probenecid, an anion pump blocker helps in retaining the thallium sensitive form of FluxOR™ reagent in the cytosol). The baseline fluorescence intensity was then measured for 2 minutes. After measuring the baseline intensity, stimulus buffer (contains thallium) was injected in each sample followed by immediate measurement of fluorescence kinetics for 4 minutes. The fluorescence intensity was measured at excitation 480 nm and emission 520 nm using a microplate reader.

### 10. *Plasmodium falciparum* growth assay and artemisinin growth inhibition assay (GIA)

*Plasmodium falciparum* laboratory strain 3D7, cultured in RPMI 1640 (Invitrogen, USA) was used for most of the *in-vitro* assays. Parasites were maintained under standard conditions. Schizonts stage infected RBCs were purified using percoll mediated density gradient for all assays*. P. falciparum* growth assay was carried out as previously described (44). In brief, purified schizonts were added to the RBCs from different ATP2B4 genotype donors, at 0.2% parasitemia and 2% hematocrit, plated in duplicate and incubated at 37 °C with 5% CO2, 5% O2 and 90% N. Initial parasitemia at 0 h and final parasitemia after 48h was measured using flow cytometer as described (45) and was also validated by microscopy. Growth rates of parasite were determined using formula: (final parasitemia—initial parasitemia)/ initial parasitemia. *P. falciparum* susceptibility to artemisinin in RBCs of different PMCA4b level was assessed (46) in the presence of range of concentrations of artemisinin from 50nM to 1.5nM. IC50 was calculated by plotting percent inhibition of growth against the drug concentration on GraphPad Prism version 8.0.1.

### 11. Growth inhibition assay by PMCA4b inhibitors resveratrol and ATA

For the growth inhibition assay, PMCA inhibitors Resveratrol and Aurintricarboxylic acid (ATA) were used. In brief ring stage *P. falciparum* were seeded in 96 well plate. Both the drugs were serially diluted and plated in duplicate. Parasitemia was monitored after 72h of incubation by counting the percentage of parasites in Giemsa-stained smears under light microscope. All experiments were performed in duplicate and 50% inhibitory concentration (IC50) was determine using dose-response curve generated in graph pad prism software.

### 12. Erythroid progenitor BEL-A cell culture

Bel-A Cells were maintained as described previously (47). Briefly, cells were cultured in expansion medium (StemSpan SFEM II (Stemcell Technologies Inc.), 50 ng/ml SCF (Immunotools, Germany), 3 U/ml EPO (Zydus, India) and 10^−6^ M dexamethasone (Sigma-Aldrich)). The cells were maintained in this medium at a density of 1–3 × 10^5^ cells/ml at 37 °C, 5% CO2. Culture medium was changed 2-3 times per week and replaced with fresh medium.

### 13. Transfection of siRNA targeting PMCA4b encoding gene ATP2B4 in BEL-A cells

ATP2B4 knockdown was done in pro-erythroblast stage of BEL-A cells, transfected with siRNA designed against ATP2B4 at two different concentration (150 nM, and 200nM). In control equal number of cells (0.5 × 10^6^) were transfected with scrambled siRNA at 1100V, 30ms, 3 pulse using Neon electroporation system (Invitrogen). After 6 h of transfection, cells were washed with 1X PBS and re-suspended in expansion medium. After 72 h of transfection, and cells were analyzed for ATP2B4 expression.

### 14. Erythrocytic topographical analysis by Atomic Force Microscopy

To examine the morphological changes in the RBCs from donors with high and low PMCA4b expression, RBCs were thin smeared and air-dried on a clean glass slide. Imaging was done using the WITec alpha system, using NSG30 probes with the force constant of 22–100 N/m, resonant frequency of 240–440 Hz, tip curvature radius of 10 nm in non-contact mode. Topographic images were obtained at a resolution of 512 ×512 points per line with the scan rate of 0.5 times/line (Trace) (s). All the AFM images were captured using the Control Four 4.1 software and analyzed using software Project Four 4.1 software (WITec, Germany).

### 15. Statistical and data analysis

To check the association of genotypes (major, minor, and heterozygous) of each regulatory regions (Enhancer, 5′UTR and 2^nd^ intron) among the four malaria outcome groups, Chi-square test for independence was used. For all the flow cytometry experiments, 20,000 events were captured, and the results were analyzed using AMNIS IDEAS software, version 6.4, to determine MFI cells following the gated selection of singlet population.

Flow cytometer data is presented as graph generated by GraphPad Prism 8.0.1. During the experiments, two control samples were included in every batch as a baseline, with their MFI values normalized to 1. The MFI of other samples was calculated relative to these controls. This normalization approach was necessary to avoid inter-batch variations in fluorescence measurements. Independent t-test was used to compare the mean between two group and one-way ANOVA was used for comparison involving more than two groups. Differences in the PMCA4b expression, intracellular calcium concentration and intracellular ROS, across different genotypes was evaluated using one-way ANOVA followed by post-hoc comparisons (Tukey’s multiple comparisons test) to compare pairwise difference between groups by excluding one genotype (Minor) as it has only one data point. Linear regression and Pearson correlation is used for all the correlation data. In all statistical analyses, a 95% confidence interval was applied to ensure the reliability of the results. Data were considered statistically significant if p < 0.05. p < 0.05 represents *, p < 0.01 represents **, and p < 0.001 represents *** significance. All analyses were performed using GraphPad Prism 8.0.1.

## Contributors

Conceptualization: PA, SG, PKM and SS; methodology: PA, SG, PM and SB; Investigation: PA, SG, PM, SB, PKM and SS; Funding acquisition: PKM and SS; Resources: PKM and SS; Sample collection: PA, VK, MPS, SKC, SS, OPS, GD, PKM; Writing, PA, SG, PKM and SS. All authors have read and approved the manuscript.

## Data sharing

The raw data supporting the conclusions of this article will be made available by the authors, on request, to any qualified researcher.

## Acknowledgments

PA is supported by CSIR through CSIR-SRF fellowship, SG is supported by DBT Indo Swiss Grant. The authors would like to express their gratitude to all the donors to participate in the study. The director ICMR-NIMR is acknowledged for providing infrastructure and research facility. The authors would like to thank AIRF for access to AFM facility at JNU. Funding from the Indian Council of Medical Research (ICMR), Indo-Department of Biotechnology (DBT) and Swiss National Science Foundation (SNSF) for SS is acknowledged.

