## Supplementary Information for "Erythrocytic plasma membrane calcium ATPase (PMCA4b) variations mediate redox imbalance to determine artemisinin sensitivity in malaria"

**Table 1: Summary of severe/mild malaria protective/susceptible [SMPA/MMSA, MMPA/MMSA] alleles of ATP2B4 gene.**

**Table 2:** **PCR condition and primers used for the amplification of regulatory DNA region of ATP2B4.**

**Table 3: Hardy Weinberg equilibrium analysis of 14 ATP2B4 SNPs.**

**Figure 1: ATP2B4 regulatory regions identified by UCSC genome browser.**

**Figure 2: Linkage Disequilibrium plot of 14 SNPs of ATP2B4 across populations of 1000 genome project.**

**Figure 3: Relationship between calcium level and SSC.**

**Appendix table 1: Summary of the previously reported severe/mild malaria protective/susceptible [SMPA/MMSA, MMPA/MMSA] alleles of ATP2B4 gene.**

This table provides a summary of malaria protective and susceptible allele based on previous research. The reference listed support the reported association. SMPA-severe malaria protective allele, SMSA- severe malaria susceptible allele, MMPA-mild malaria protective allele, MMSA- mild malaria susceptible allele.

| **SNP ID** | **Significance from previous study** | **Reference** | **Alt allele frequency in Indian database** |
| --- | --- | --- | --- |
| rs10900585 (G>T) | G- associated with decreased risk of SM. (African population) **(SMPA)**  G- reduced malaria anaemia, protect against malaria in pregnancy. (African population) | 2012-Timmann et.al.  2013- Bedu-Addo et.al. | 0.8832 |
| rs10751450 (C>T)  rs10751451 (C>T)  rs10751452 (T>C) | Associated with PMCA expression, RBC traits, malaria susceptibility (multicentre study, African population)  Risk genotype for severe malaria and mild malaria in Sangelese population -CC, CC, TT (African population) **(SMSA)** | 2014-Malaria Genomic Epidemiology Network  2017- Lessard et.al.  2021-Mozner et.al  2022-Nisar et.al.  2023-Thiam et.al. | 0.8857  0.8803  0.8802 |
| rs1541252 (T>C)  rs1541253 (T>C)  rs1541254 (C>G) | Risk genotype for severe malaria and mild malaria - CC, CC, GG (African population) **(SMSA, MMSA)**  TT, TT, CC- low PMCA in healthy volunteers in Hungary. (European population)  TT-Reduced PMCA expression, slow growth of Pf. (Gambia) | 2014-Nadila et.at.  2017-ambo et.al.  2022-Nisar et.al.  2022-Joof et.al. | 0.8802  0.8743  0.8834 |
| rs15411255 (G>A) | GG- reduced parasite density. | 2022-Uyoga et.al. | 0.8743 |
|  | PMCA inhibitor reduced malaria parasite growth. | 2022-Puji et.al. |  |

**Appendix table 2: PCR condition and primers used for the amplification of regulatory DNA region of ATP2B4.**

| **Gene/ region** | **Primers** | **Amplicon size** | **1° PCR** | **2° PCR** |
| --- | --- | --- | --- | --- |
| ATP2B4 (Enhancer) | 5’AGAGGATTGTCAGGAATCGGC3’  5’TCCAGGTGTATGGAAATGACAAC3’ | 887bp | 95°C-3min  95°C-30sec  55°C-1 min  72°C-4 min  72°c-10min | 95°C-5min  95°C-30 sec  63°C-30sec  72°C-1 min  72°C-5 min |
| ATP2B4 (5’UTR) | 5’ACTGGGTACCTCTTGCCCTT3’  5’CACTAGCCACCTTCCTCCAA3’ | 749bp |  |  |
| ATP2B4 (2^nd^ intron) | 5’TGGGATGCTAAATCACAAGGT3’  5’GGCCAAGTAAGAATCTGCTACA3’ | 603bp |  |  |

**Appendix table 3: Hardy Weinberg equilibrium analysis of 14 ATP2B4 SNPs.**

| **SNPs** | **chi-square test statics** | **P-value of chi-square** |
| --- | --- | --- |
| rs11240733 | 0.00012013 | 0.991255051 |
| rs10751449 | 0.00729955 | 0.931913657 |
| rs10736845 | 0.00729955 | 0.931913657 |
| rs10751450 | 0.051870221 | 0.81984033 |
| rs10751451 | 7.12142E-08 | 0.999787077 |
| rs10751452 | 0.003703729 | 0.951472087 |
| rs1541252 | 0.0013571 | 0.970613504 |
| rs1541253 | 0.0013571 | 0.970613504 |
| rs1541254 | 0.004161543 | 0.94856413 |
| rs1541255 | 0.004161543 | 0.94856413 |
| rs1419114 | 0.00068943 | 0.979052353 |
| rs10900585 | 0.175234726 | 0.675500836 |
| rs10900586 | 0.015020412 | 0.90245728 |
| rs10793762 | 0.01115214 | 0.915896793 |

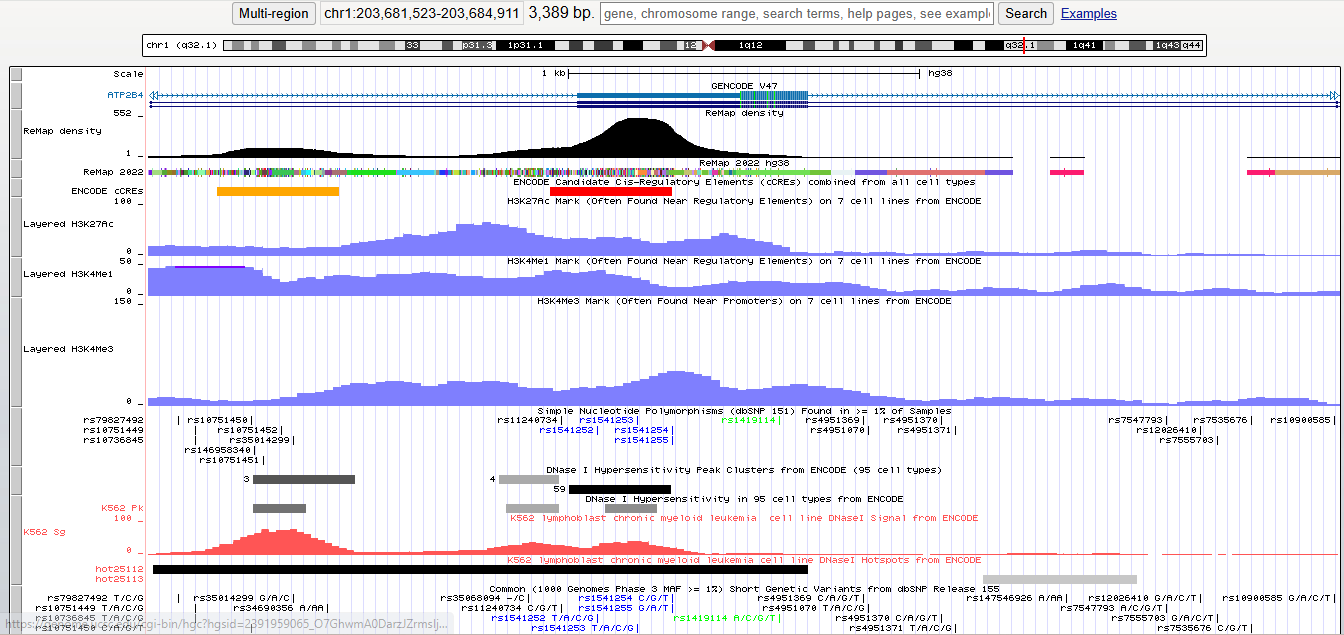

**Appendix figure 1:** **ATP2B4 regulatory regions identified by UCSC genome browser.** ATP2B4 region ch1:203681464-203684970 containing the 8 variants studied for regulatory mechanism. **[A]** Remap tool is a computational platform used for analysing and interpreting functional genomics data. It integrates various publicly available ChIP-seq (Chromatin Immunoprecipitation sequencing) datasets to identify regulatory elements such as enhancers, promoters, and transcription factor binding sites across the human genome. Out of 3 regions, enhancer and 5’UTR region are located within main peaks of Remap density. Candidate Cis-Regulatory Elements [cCREs] in the human genome are displayed. Red colour indicates cCREs with promoter-like signatures [cCRE-PLS] in 5’UTR region and orange colour indicate proximal cCREs enhancer-like signatures [cCRE-ELS] in enhancer region. H3K27Ac (acetylation of lysine 27 of the H3 histone protein) and H3K4me1 (tri-methylation of lysine 4 of the H3 histone protein) marks are associated with enhancers showing peaks in enhancer and 5’UTR, while H3K4me3 is associated with active promoters showing main peak in 5’UTR. DNase I hypersensitivity regions display peaks at enhancer and 5’UTR sites, indicating their regulatory role in ATP2B4 gene transcription.

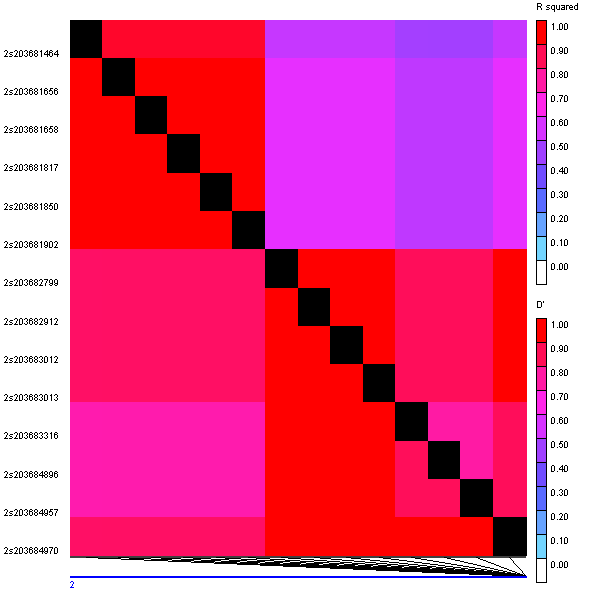

**Appendix** **figure 2: Linkage Disequilibrium plot of 14 SNPs of ATP2B4 across populations of 1000 genome project.** The LD (Linkage Disequilibrium) plot showcases the patterns of pairwise linkage disequilibrium, measured by R-squared (R^2) and D' statistics, among 14 SNPs situated in the regulatory regions of ATP2B4 gene variants within all population of 1000genome project. The plot employs color-coded representation to signify the strength of LD between SNP pairs.

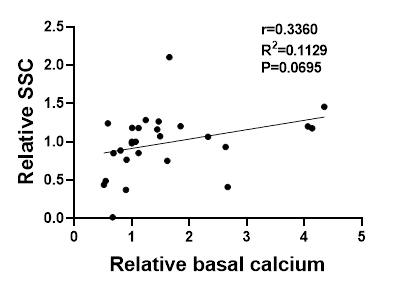

**Appendix figure 3: Relationship between calcium level and SSC.** Scatter plot depicting the relationship between basal calcium levels and relative side scatter of RBCs. A weak positive correlation was observed.
